# Serotonin signaling modulates aging-associated metabolic network integrity in response to nutrient choice

**DOI:** 10.1101/2021.03.01.433267

**Authors:** Yang Lyu, Daniel E.L. Promislow, Scott D. Pletcher

## Abstract

Aging arises from complex interactions among multiple biochemical and metabolic products. Systems-level analyses of biological networks may provide insights into the causes and consequences of aging that evade single-gene or single-pathway studies. We have shown that dietary choice *per se* is sufficient to modulate aging and metabolic health in the vinegar fly, *Drosophila melanogaster*. In other words, how each meal is presented, or the way in which it is eaten, is influential, independent of the amount or type of nutrients that are consumed. For example, when major macronutrients were presented separately, male flies exhibited a rapid and significant increase in mortality rate and a reduced overall lifespan relative to those fed a single medium containing both sugar and yeast. These effects are mediated by specific components of serotonin signaling, as a mutation in serotonin receptor 2A (*5-HT2A*) eliminated the effects of dietary choice. Here we show that dietary choice influenced several measures of metabolic network integrity, including connectivity, average shortest distance, community structure, and robustness, with the effects of the latter two restricted to tissues in the head. These changes in network structure were associated with organism resilience and increased susceptibility to genetic perturbation, as measured by starvation survival. Our data suggest that the behavioral or perceptual consequences of exposure to individual macronutrients, involving serotonin signaling through 5-HT2A, qualitatively change the state of metabolic networks throughout the organism from one that is highly connected and robust to one that is fragmented, fragile, and vulnerable to perturbations.

## Introduction

Emergent, system-level properties of biological networks have provided insights into many complex behaviors of organisms that single-gene or pathway analyses often struggle to explain^1–3^. For example, robust gene networks allow animals to maintain segment patterning against perturbations during embryonic development, through interactions and feedback controls among multiple transcriptional modules within and between cells^4^. Regulatory loops in signal transduction networks render bacterial chemotaxis immune to intrinsic noises^5^. During eye development in *Drosophila melanogaster*, *microRNA-7*, transcriptional factor *Yan*, and their downstream targets together form a feedback system to stabilize photoreceptor determination processes against temperature fluctuations^6^. These examples illustrate that a network perspective on how molecules function together in biological processes provides us with valuable insight into underlying molecular mechanisms.

Here we explore the role of metabolic network properties in the context of biological aging. Aging arises from complex interactions among multiple biochemical and metabolic products^7–9^, and these networks may decline in integrity with advancing age^10^. In the aged tissues of mice, for example, expression variation in single genes is significantly increased, and expression correlation between genes is decreased, relative to similar measures from young animals^11, 12^. A causal link between longevity and network integrity has been suggested by data from fruit flies and nematode worms showing that specific anti-aging interventions, such as dietary restriction, promote the connectivity of transcriptomes and metabolomes ^13,14^. Mechanistically, aging may diminish network integrity by preferentially affecting the ‘hubs’, because molecules that have more partners^15–17^ or that connect to multiple functional groups^18^ are more likely to influence lifespan. Interestingly, metabolic network structures appear to be dynamically regulated - hub positions shift in response to environmental conditions (e.g., nutrient availability^19^), although the mechanisms responsible are largely unknown. Sensory perception is likely a key initiator of such changes, while the consequences may be substantial for integrative traits, such as health and lifespan.

Neuroendocrine systems are important mediators linking sensory perception, lifespan, and metabolic health^20, 21^. In humans, serotonin signaling coordinates a range of behavioral and physiological traits including emotion, sleep, feeding and metabolism (e.g. Refs ^22, 23^). These regulatory mechanisms are well conserved across taxa, with many new insights first described in invertebrate systems such as *Caenorhabditis elegans*^24–26^ and *Drosophila*^27–29^. For instance, research from *C. elegans* revealed that, despite their well-known role in modulating appetite and feeding behavior^22, 30^, specific serotonin receptors regulate fat metabolism independent of feeding^25^. Most recently, we discovered that serotonin 2A (5-HT2A) receptor in *D. melanogaster* modulates lipid storage, stress resistance, and longevity in response to nutrient availability and independent of feeding^31^.

While investigating how 5-HT2A is involved in coordinating dietary conditions with physiology and aging, we found that it was required for diet-dependent changes in the abundances of metabolites from central metabolic processes^31^. These included TCA cycle intermediates and their amino acid precursors, which govern energy homeostasis and nutrient flux through many metabolic networks. We therefore speculated that changes in these hub metabolites may affect the hidden structure of broader metabolic networks and that such changes in network structures *per se* may modulate lifespan^10, 13^. Indeed, we found that diet- dependent changes in many characteristics of network integrity, including measures of connectivity, average shortest distance, module clustering, and robustness are mediated by serotonin signaling through the 5-HT2A receptor. Our findings therefore ascribe a role for this ancient signaling pathway in regulating systems-levels properties and, to our knowledge, provide the first demonstration of a molecular mechanism responsible for changes in metabolic network structure that may modulate aging.

## Results

### Dietary choice decreases metabolite co-expression through 5-HT2A

Previously, we demonstrated that 5-HT2A modulates aging in *Drosophila* in response to how macronutrients are presented to the animals^31, 32^. We found that dietary choice (i.e. presenting 10% w/v sugar and 10% w/v yeast in separate wells) increased mortality (Figure 1A) and reduced mean lifespan of wildtype males when compared with siblings that were presented with a single, complete food of the same macronutrient composition (i.e. 10% w/v sugar and 10% w/v yeast mixture)^31^. A dietary switch at 20 days of adult life rapidly increased/decreased mortality when flies were transferred to a choice or fixed diet, respectively (Figure 1B and 1C). Reduced expression of *5-HT2A*, caused by a transposable element insertion in its promoter region^33^ eliminated the differences in mortality between the two dietary environments (Figure 1D). 5-HT2A was also required for the effects of nutrient presentation on the metabolome: loss of wild-type *5-HT2A* gene function diminished the changes in metabolite abundances caused by dietary choice, where critical TCA intermediates and their amino acid precursors were significantly increased in wildtype animals when they had to choose between sugar and yeast before each meal ^31^. The questions that we address herein are whether dietary choice, and serotonin signaling through this receptor, also affect metabolic network structures and whether these system-level changes influence health and aging.

**Figure 1.**
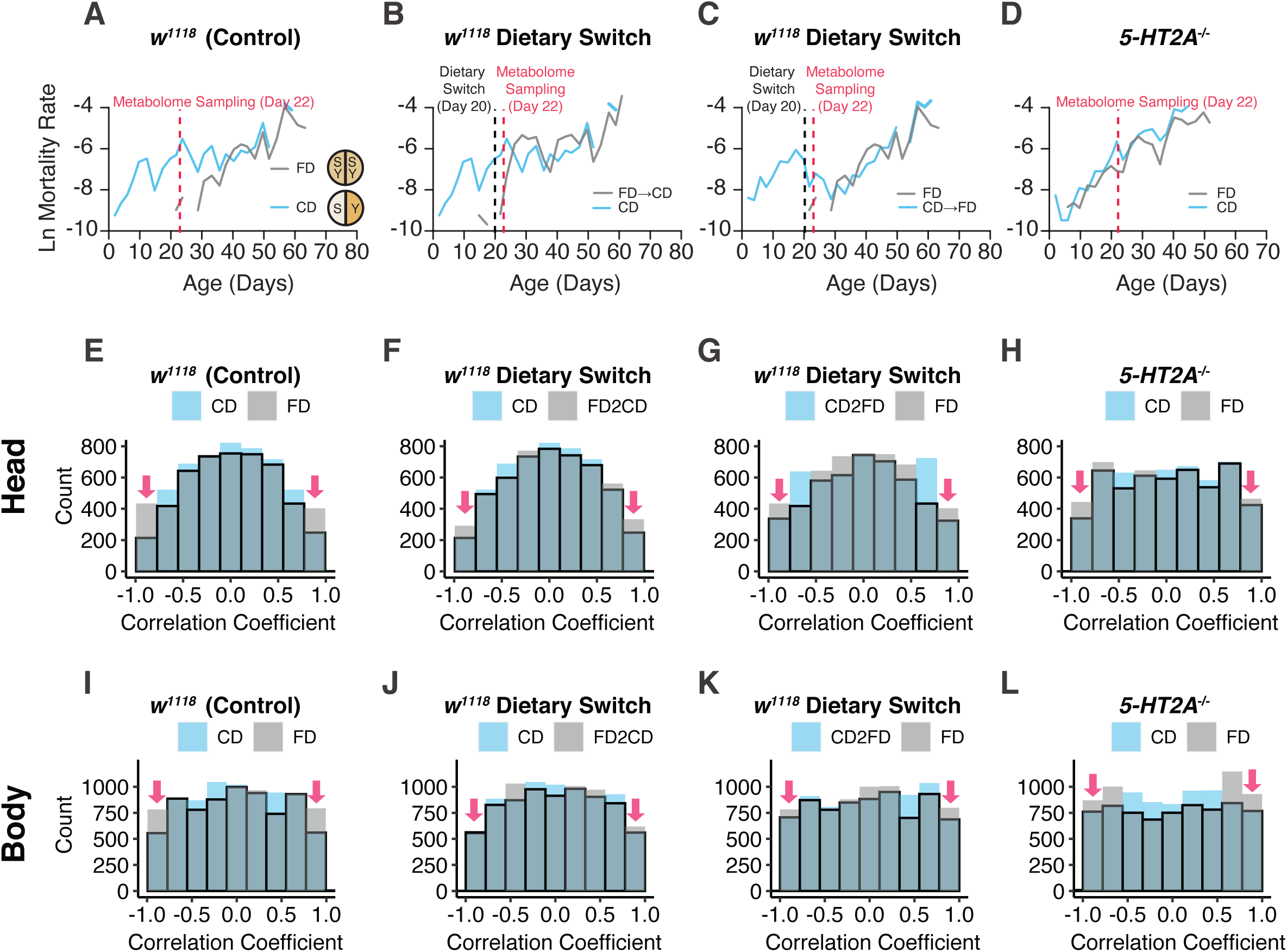
Dietary choice influences both mortality and metabolite correlations in metabolomes through serotonin signaling. (**A-D**) Mortality data showing the choice diet effects, which are from our recent study (Lyu et al. 2021) and used to demonstrate the association between aging and metabolite correlations. Mortality rate is plotted on the natural log scale (N = 225-282). (**A**) Control males fed on a choice diet exhibits higher mortality rates early compared to their siblings on a fixed diet. (**B-C**) Switching to a different diet (applied at 20 days of adulthood, as denoted by the dashed line) immediately changes the mortality rates that reflects the influences from the new diet. (**D**) *5-HT2A* mutants are immune to the choice-induced mortality effects. (**E-L**) Dietary choice impacts correlations between metabolites in both heads and bodies. (**E and I**) Control males fed a choice diet have less highly correlated metabolite pairs (indicated by red arrows) compared to their siblings that fed a fixed diet. Either dietary switches (**F, G, J, and K**) or mutation in *5-HT2A* (**H and L**) diminish the dietary differences, mirroring the effects on mortality.

Targeted metabolomics data were collected from the heads and bodies of 22-day-old adult flies that had been provided either a choice diet (CD) or a fixed diet (FD) since two days post- eclosion. We also examined flies that were maintained on these diets until 20 days of age and then switched two days prior to sampling, either from a choice diet to one that is fixed (CD &#x2192;FD) or *vice versa* (FD CD). The age at which flies were switched was chosen as that time when the mortality differences between the CD and FD cohorts were maximized (Figure 1A). We began our interrogation of diet-dependent metabolic networks by calculating correlations between all pairs of metabolites across replicate samples within each diet treatment. We chose this approach because changes in physical networks (e.g., gene regulatory and metabolic networks) are often reflected in changes in the correlations between pairs of transcripts or metabolites^13^, and previous studies have suggested that correlation coefficients (*ρ*) between molecules might decrease with advancing aging in fruit flies and mice ^12, 13^.

We found that the correlations between metabolites in flies fed on a choice diet were significantly weaker than those from flies fed a fixed diet (Figure 1E). The percentage of highly correlated metabolite pairs (i.e., those with |*ρ*| ≥ 0.8) in the heads of flies fed the choice diet was 7.7%, compared to 13.9% from flies fed the fixed diet (*P* < 0.001, Fisher’s exact test). A similar trend was observed in bodies (Figure 1I), with a 28.2% reduction of highly correlated pairs in flies given a dietary choice (12.2% vs 17.0%, flies on choice vs. fixed diet, respectively; *P* <0.001, Fisher’s exact test). The differences were reduced upon dietary switch (Figure 1F, 1G, 1J and 1K). These data reveal an effect on metabolite correlation structure in control animals that is associated with dietary choice and that is temporally coincident with changes in age- specific mortality.

Disrupting serotonin receptor *2A* abrogated the differences in mortality between diets (Figure 1D), which led us to ask whether the effect of diet on metabolite correlation structure was also mediated through the 5-HT2A receptor. Indeed, we found that the loss of highly correlated metabolite pairs in the choice diet environment was significantly reduced in mutant animals by roughly 39% in the heads (from a 6.2% decrease in control animals to 3.8% in *5-HT2A* mutants, *P* < 0.001, Fisher’s exact test) and 19% in the bodies (from a 4.8% decrease by choice in the control genotype to 3.9% in mutants, *P* = 0.030, Fisher’s exact test). Our observations suggest that the way in which nutrients are presented influences the physical nature of the metabolome and that these changes in network structure are mediated, at least in part, by serotonin signaling.

To better understand the extent of changes in network structure induced by our dietary manipulation, we used adjacency matrices to construct correlation networks among metabolites. We considered two metabolites to be directly linked if they were significantly correlated with each other (significance was estimated using Spearman’s rank-order correlation followed by FDR correction). These connections formed the edges of our metabolite correlation networks that link individual metabolites or nodes (see Figure 2A and Methods for more details on network construction). Biologically speaking, if two metabolites are connected by an edge, then they are thought either to interact directly (e.g., by co-involvement in the same biochemical reaction) or to share patterns of abundance, perhaps as a result of co- regulation by common mechanisms. Using this approach, we constructed one network for each combination of diet, genotype, and tissue sample. We used a threshold FDR = 0.10 to identify significant comparisons among different conditions, which is equivalent to *ρ* = 0.7-0.8 across different conditions. A more stringent criteria of FDR = 0.05 (*ρ* = 0.8-0.9 across different conditions) yielded similar results but led to a restrictive number of edges in the head network derived from control flies on a choice diet (Supplementary Figure 1). We found that networks constructed from each of the eight groups exhibited a similar organization across diets, tissues, and genotypes (see Figure 2B for networks constructed from heads and Supplementary Figure 1 from bodies). Each consisted of one large, highly connected core group comprised of at least 70% of all the observed metabolites, together with many small satellite groups, each consisting of no more than five inter-connected nodes, which are not connected with the core. Most of these satellites were comprised of a single, unconnected metabolite.

**Figure 2.**
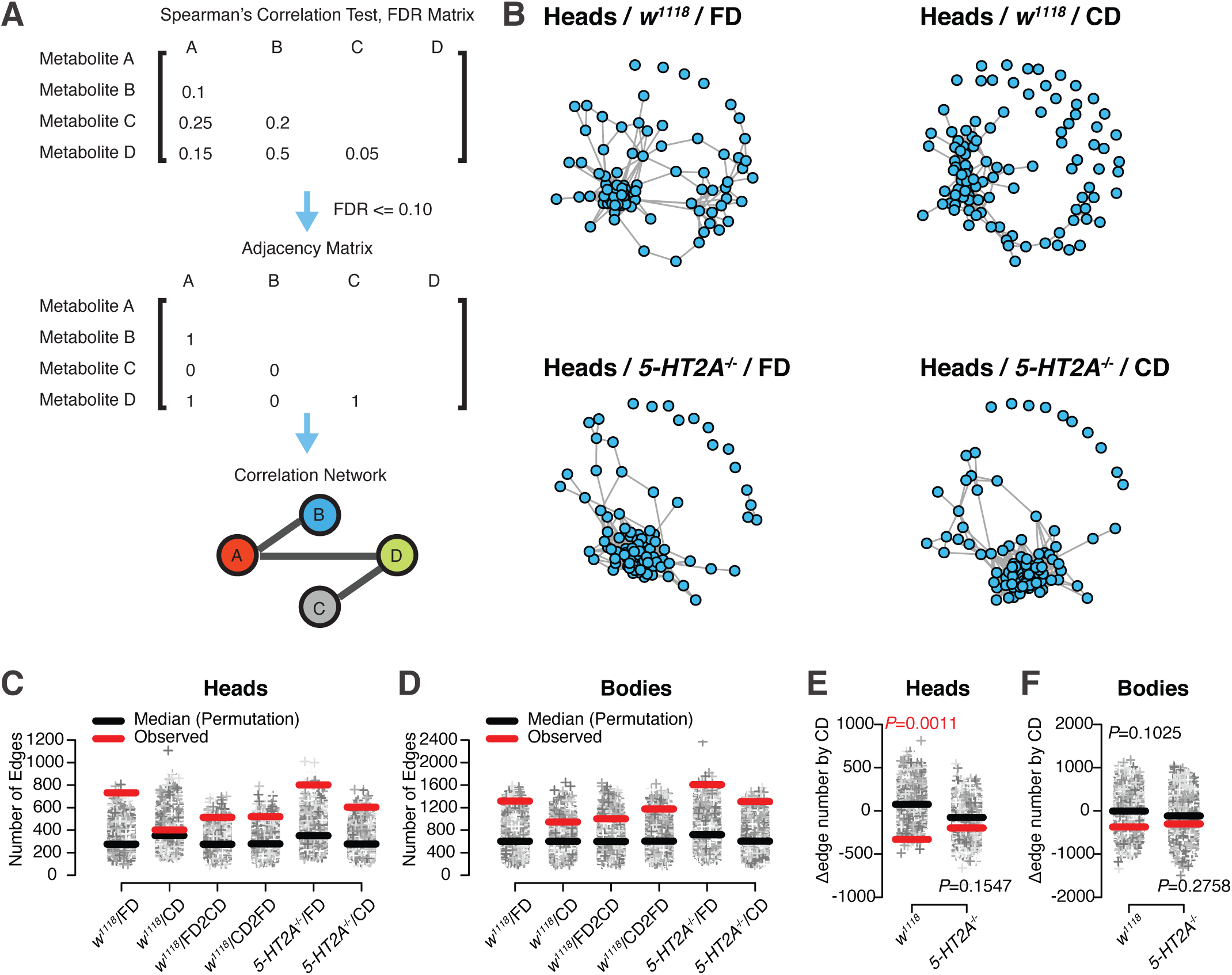
Serotonin signaling modulates the choice effects on the edge number of correlation networks. (**A**) Diagram illustrating our approach to construct correlation networks from Spearman’s rank correlation test. (**B**) Visualization of the correlation networks demonstrates similar organizations across different diets and genotypes. (**C-D**) Dot plots showing the edge numbers of the real (denoted as the red bar) and the randomized networks. Grey dots represent 10,000 simulations for each group with the black bars indicating the median. Deviations from random datasets are observed and their trends are similar between heads (**C**) and bodies (**D**). (**E-F**) Dietary differences in control males are significant in the heads (E) and the *P*-value is marginal in bodies (**F**). Such effects are not seen in *5-HT2A* mutants, suggesting the requirement of serotonin signaling.

To understand the composition of the core group, we queried it for metabolic pathway enrichment. A complete enrichment analysis was constrained by the size of our metabolomic panel, so we instead chose to highlight pathways with more than six metabolites (as shown in Table 1). We found that three major metabolic pathways, including aminoacyl-tRNA biosynthesis, purine metabolism, and glycine, serine and threonine metabolism, were highly represented in all core groups. Four pathways (alanine, aspartate and glutamate metabolism, butanoate metabolism, arginine biosynthesis and glyoxylate and dicarboxylate metabolism) that appeared in the controls fed on a fixed diet were not highly represented in the head of the same genotype that fed on a choice diet. Two pathways (butanoate metabolism as well as alanine, aspartate and glutamate metabolism), however, were highly represented in the heads of 5-HT2A mutant flies fed a choice diet.

Pathway analysis on the metabolites from each core group. The number of metabolites are shown only if they are greater than six.

**Table 1. Pathway analysis on the metabolites from each core group. The number of metabolites are shown only if they are greater than six.**
|  | # of Metabolites in the Pathway (Heads) |  |  |  | # of Metabolites in the Pathway (Bodies) |  |  |  |
| --- | --- | --- | --- | --- | --- | --- | --- | --- |
|  | *w*<sup>1118</sup> |  | *5-HT2A*<sup>-/-</sup> |  | *w*<sup>1118</sup> |  | *5-HT2A*<sup>-/-</sup> |  |
|  | FD | CD | FD | CD | FD | CD | FD | CD |
| Aminoacyl-tRNA biosynthesis | 20 | 16 | 18 | 17 | 19 | 19 | 19 | 18 |
| Purine metabolism | 11 | 11 | 11 | 13 | 12 | 12 | 13 | 13 |
| Alanine, aspartate and glutamate metabolism | 10 |  | 10 | 10 | 11 | 11 | 11 | 10 |
| Glycine, serine and threonine metabolism | 9 | 7 | 9 | 9 | 9 | 9 | 9 | 9 |
| Butanoate metabolism | 7 |  |  | 7 | 7 | 7 | 7 |  |
| Arginine biosynthesis | 7 |  |  |  | 8 | 8 | 7 | 8 |
| Glyoxylate and dicarboxylate metabolism | 7 |  | 7 |  | 7 | 7 | 7 | 7 |

### Serotonin signaling modulates edge number, connectivity, and average shortest distance of metabolite correlation networks

We next asked whether the finer structures of the networks were influenced by dietary presentation. We began by focusing on edge number because of our results that revealed an influence of diet on correlation coefficients (Figure 1E-1L). Permutation analysis was used to compare the edge number of real networks with that of randomly shuffled ones (n=10,000, see Methods for details) and to identify significant edge differences among dietary conditions. In the observed network derived from the heads of *w^11^*^18^ flies fed a fixed diet, the number of edges placed the network in the upper extreme of the distribution of edges drawn from randomized networks (*P* < 0.002; red bar vs black bars in the first column of Figure 2C), while for the observed network drawn from heads of flies maintained on a choice diet, the number of edges was not unusually high (*P* < 0.341, shown in the second column of Figure 2C). Differences of this nature were also observed in the networks constructed from body samples from the same cohorts: the observed edge number in the fixed diet network was more extreme (*P* < 0.002, as shown in the 1st column of Figure 2D) than was edge number from the choice diet group (*P* < 0.077, the second column of Figure 2D). To determine whether these differences were more than expected by chance, we compared the differences of edge number (#edge of CD - #edge of FD) between 10,000 pairs of randomly shuffled networks within each genotype and tissue. We found that the reduction in the number of edges induced by dietary choice was significant in the network constructed from the heads of control animals (*P* = 0.0011, permutation test, the first column in Figure 2E). A similar trend was observed in the metabolic networks constructed from fly bodies, but the significance of diet was marginal (*P* = 0.1025, permutation test, the first column in Figure 2F). If reduced edge number is associated with increased mortality and faster aging, then we would expect that manipulations that modulate mortality should also affect edge number. Indeed, dietary switches, which led to rapid changes in mortality, also stimulated a 48hr modification of metabolic network edge number in the direction that reflected the influence of the new diet. These changes were observed in both heads and bodies (depicted as the 3rd and 4th columns in Figure 2C and 2D), and they mirror the changes in mortality.

We investigated whether receptor 5-HT2A was required for the reduction in edge number induced by dietary choice. We found that the diet effect was reduced in *5-HT2A* mutant flies (Figures 2C and 2D) compared to control animals, primarily due to more edges in the network constructed from flies on a choice diet. In mutant flies, edge differences between diets were not statistically significant (*P* = 0.15 in heads and *P* = 0.28 in bodies, Figure 2E and 2F), indicating that these effects are 5-HT2A dependent.

This led us to examine the distribution of the number of edges connected to each metabolite, which is referred to as the degree, in our correlation networks. Metabolite degree exhibited a bimodal distribution under most conditions in both heads (Figure 3A) and bodies (Figure 3B), where low-degree and high-degree nodes were clearly overrepresented. Notably, we observed a strong reduction in high-degree nodes in the heads of control flies fed a choice diet but no effect of diet on degree distribution in 5-HT2A mutants diet (Figure 3A, compare right two plots). We defined high-degree nodes relative to core size (i.e., those with degree ≥ 30% × total node number), to exclude the possibility of confounding between the two (Figure 2B). High- degree nodes accounted for 40.5-59.0% of the total metabolites in all conditions (indicated as red dots in Figure 3A and 3B), except for measures from the heads of control flies fed a choice diet, where none of the nodes met the criteria. The loss of high-degree nodes was not observed in the bodies of the same cohort, nor in the heads of 5-HT2A mutants (40.5% and 50.5% of the metabolites were accounted as high-degree, respectively, also see Figure 3B). This suggested that serotonin signaling reduced neural network connectivity by removing ‘hub’ genes in response to dietary choice^34–36^.

**Figure 3.**
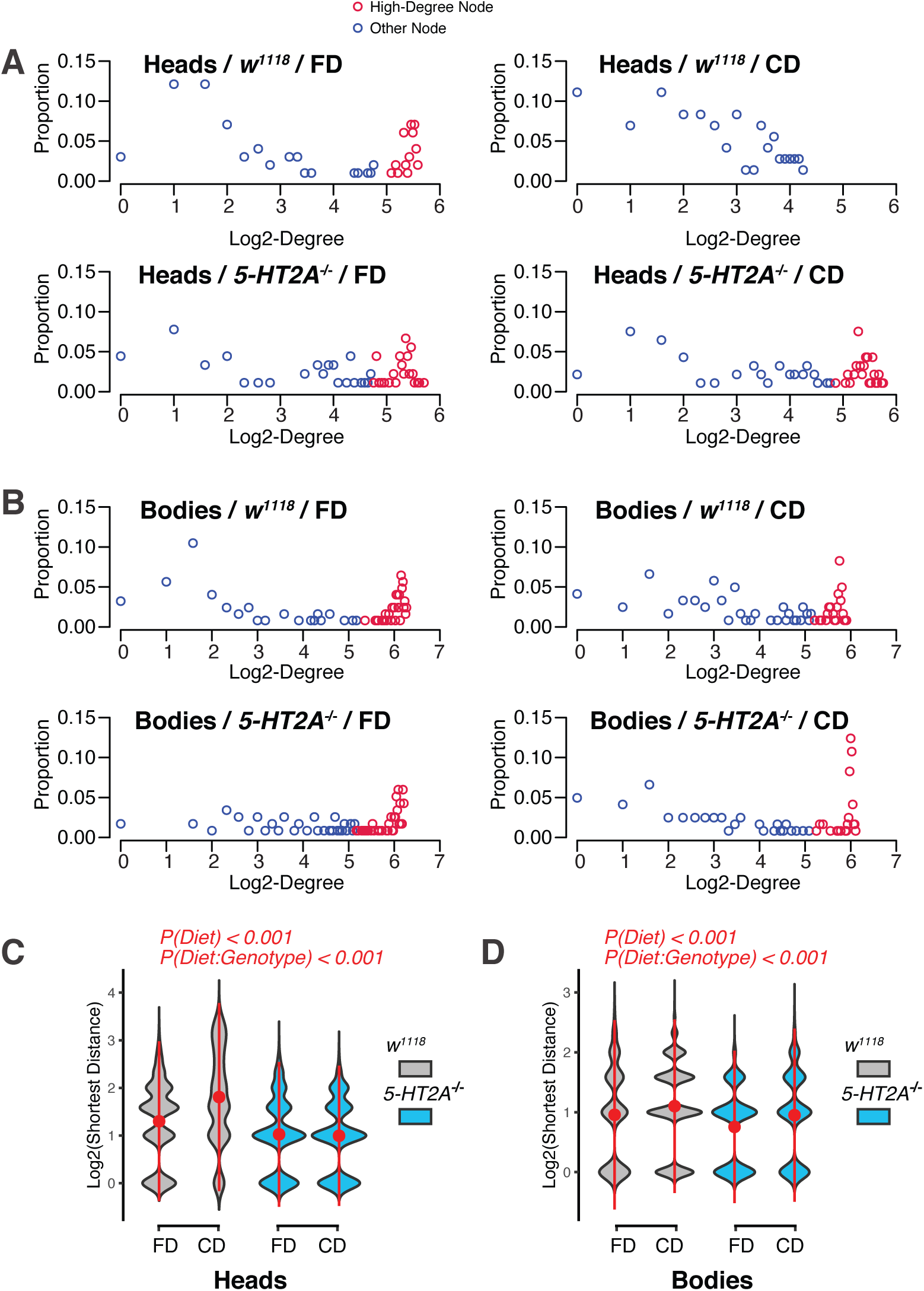
Choice environment reduces network connectivity and increases average shortest distance in a 5-HT2A dependent manner. (**A-B**) Dot plots on the frequencies of node connectivity (degree) demonstrate decreases in the number of high-degree nodes is in the heads (**A**), but not the in the bodies (**B**) of control flies that fed on a choice diet (compared to that of flies fed a fixed diet). On the other hand, node connectivity of *5-HT2A* mutants are not influenced much by diets. (**C-D**) Exposure to a choice diet also increases the average of shortest distance in networks and such effects are dependent on *5-HT2A* in both heads (**C**) and bodies (**D**). *P*-values on top of the violin plots are obtained from Two-way ANOVA.

Because random pairs of metabolites are most likely to be connected through network hubs, we conjectured that the average path length between two metabolites in the network would also be affected by nutrient presentation. If true, we wanted to determine whether this effect was mediated by *5-HT2A*. We measured the average of the shortest path length (see Methods for definition) across all metabolite pairs^37^. Generally, having fewer edges in the network results in fewer possible routes between two metabolites, thus generally increasing the distance between them, but this is not necessarily so. We found that the average shortest distance (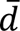) between metabolites was significantly increased in networks calculated from the heads and bodies of control flies in a dietary choice environment compared to the corresponding samples from the fixed diet (Figure 3C and 3D, One-way ANOVA, *P*(Diet) < 1.1 for control flies). These changes were dependent on *5-HT2A* in both tissues (two-way ANOVA, *P*(Diet:Genotype) < 0.001). Interestingly, the values of 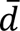 in these networks (ranging from 1.9 to 4.4) were small relative to the size of the networks (network diameters, which are the longest paths in the networks, range from 5-13). The relationship is consistent with small- world architecture^38, 39^, documented in many previously examined metabolic networks^34–36^.

### Dietary choice induces the fragmentation of the network in a 5-HT2A dependent manner

Biological networks are often organized into tightly-knit modules, within which metabolite- metabolite interactions are dense and between which there are only looser connections^40^. Here, in metabolite co-expression networks, modules might represent a group of functionally related metabolites that respond to dietary inputs in a highly coordinated manner. We investigated dietary influence on the organizational structure of metabolites by comparing the modules from animals that fed on different diets. Metabolite modules were determined using Newman’s leading eigenvector algorithm41, which is built around the concept that the optimal metabolite modules are those with maximal “modularity score” (see Materials and Methods for details). We then visualized the modules by plotting the correlation between every pair of metabolites. Once metabolites were ranked by the group order, modules were represented as visually perceivable blocks (e.g., Figures 4A and 4B).

**Figure 4.**
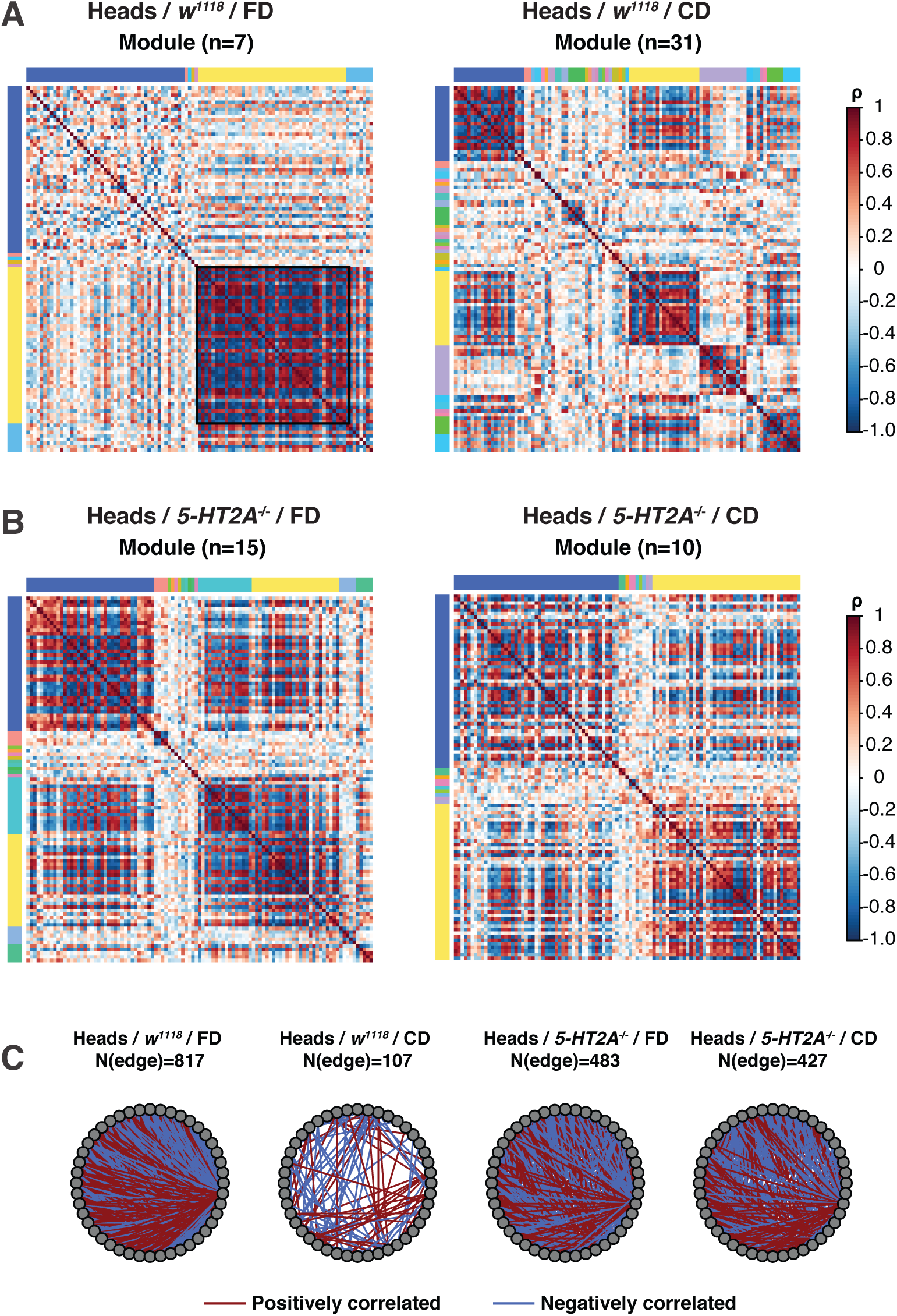
Dietary choice induces the fragmentation of network partially through 5-HT2A. (**A-B**) Correlation plots showing the dietary influences on the community structures in both control males (**A**) and 5-HT2A mutants (**B**). In controls that fed on a choice diet, the number of modules is increased, while the size of the dominant module (labeled as yellow bar) is decreased in metabolomes. Such dietary differences are eliminated in *5-HT2A* mutants. (**C**) Metabolite-metabolite interactions in the dominant module are largely lost in the CD networks that derived from the heads of control males, while part of them are restored in the *5-HT2A* mutants.

We found that dietary choice induced network fragmentation in heads (Figure 4A). A greater number of smaller modules were identified in the networks constructed from the heads of flies given a choice diet compared to those given a fixed diet. The two dominant modules from flies given a dietary choice were comprised of only 21 metabolites each, while the corresponding modules from fixed diet networks contained 47 and 44 metabolites, respectively. This pattern was not observed in bodies (Supplemental Figure 2). We speculated that the highly intra- connected dominant module (indicated by the yellow bar in Figure 3A, fixed diet) represented a homeostatic central unit, wherein metabolites respond to environmental input in a coordinated fashion. We examined the functional enrichment of this unit using MetaboAnalystR ^42^. Using our threshold of N > 6 metabolites (only 10-27% of the total functional categories fit this criterion), we found that purine metabolism (N = 9) and aminoacyl-tRNA biosynthesis (N = 8) were highly represented in the dominant module in fly heads. Our results suggest that central metabolic processes are less coordinated in a heterogeneous nutritional environment in which animals must choose their meal than they arein a more homogenous one.

We next asked whether the observed network fragmentation that was induced by our dietary manipulation required serotonin signaling through the *5-HT2A* receptor. Correlation plots revealed that dietary influences on the number and size of modules were largely abrogated in the heads of *5-HT2A* mutants (Figure 4B). To further quantify the influence of 5-HT2A function on the dominant module, we calculated the number of metabolite interactions (i.e., network edges) that were lost in control animals but maintained in mutant flies when both were placed on the choice diet. Interactions between metabolites inside of the dominant module were reduced by 86.9% by dietary choice (Figures 4C), while the effect of dietary choice on edge number was much smaller in networks from *5-HT2A* mutant flies (reduced by only 11.6%, *P* < 0.001, Fisher’s exact test), which suggests that 5-HT2A is partially required for reduced modularity following dietary choice. Of the 717 connections that were lost in the heads of control flies when they were switched to a choice diet (see Figures 4C), 317 were restored in *5- HT2A* mutants. For those that were restored, 79.8% (253/317) were restored in the same direction (either positive or negative) as originally observed in networks from flies fed a fixed diet. Together these data suggest that serotonin signaling affects both the size of the central metabolic unit as well as the number of modules. Specifically, 5-HT2A signaling appears to mediate the loss of metabolite interactions in the central module that affects amino acid metabolism and energy homeostasis upon dietary choice, which may be influential on mortality (Figure 1A-1C).

### Dietary choice decreases network robustness through serotonin signaling

A less connected and more fragmented network might be considered less robust and less resilient to perturbation. We examined this issue computationally, using two approaches to study the effect of node removal on the integrity of the network core. First, we measured the effect of random node removal on the average shortest distance (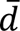) of each network core, as 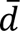 reflects how well metabolites are connected with each other in networks. It has been noted that biological networks are generally robust to random errors^43^, and indeed, we found that removal of up to 30% of the nodes (e.g., 20 nodes from the networks constructed from head and 30 nodes from body samples) had no significant effect on 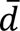 in most conditions. (Figure 5A and 5B). The only influence is in the head of control flies that fed on a choice diet, where node removal decreased 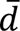 (indicated as black triangles in Figure 5A). Our second computational approach for studying robustness involved targeted network “attack”^43, 44^, in which node removal was prioritized based on measures of node importance, or ‘centrality’. We used two measures of node centrality, degree and eigenvector, which yielded similar results (see Materials and Methods). First, we observed a consistent increase in 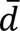 when nodes with the highest centrality were removed first and then subsequent nodes were removed in descending order (Figures 5C and 5D for degree centrality; 5E and 5F for eigenvector). Second, attacking nodes in the metabolomic networks measured from the heads of flies fed a choice diet resulted in several rises in 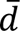, with the removing order based on either degree (Figure 5C and 5D) or eigenvector (Figure 5E and 5F). Two metabolites, 5-aminopentanoic acid and choline, were influential in both attacking methods, while GMP, isovalerate, carnitine, and glutarate were specific to one method (Figure 5C). Our data suggest that dietary choice may act through serotonin signaling to reshape the connections between metabolites in such a way as to produce a network that is relatively more vulnerable to node removal.

**Figure 5.**
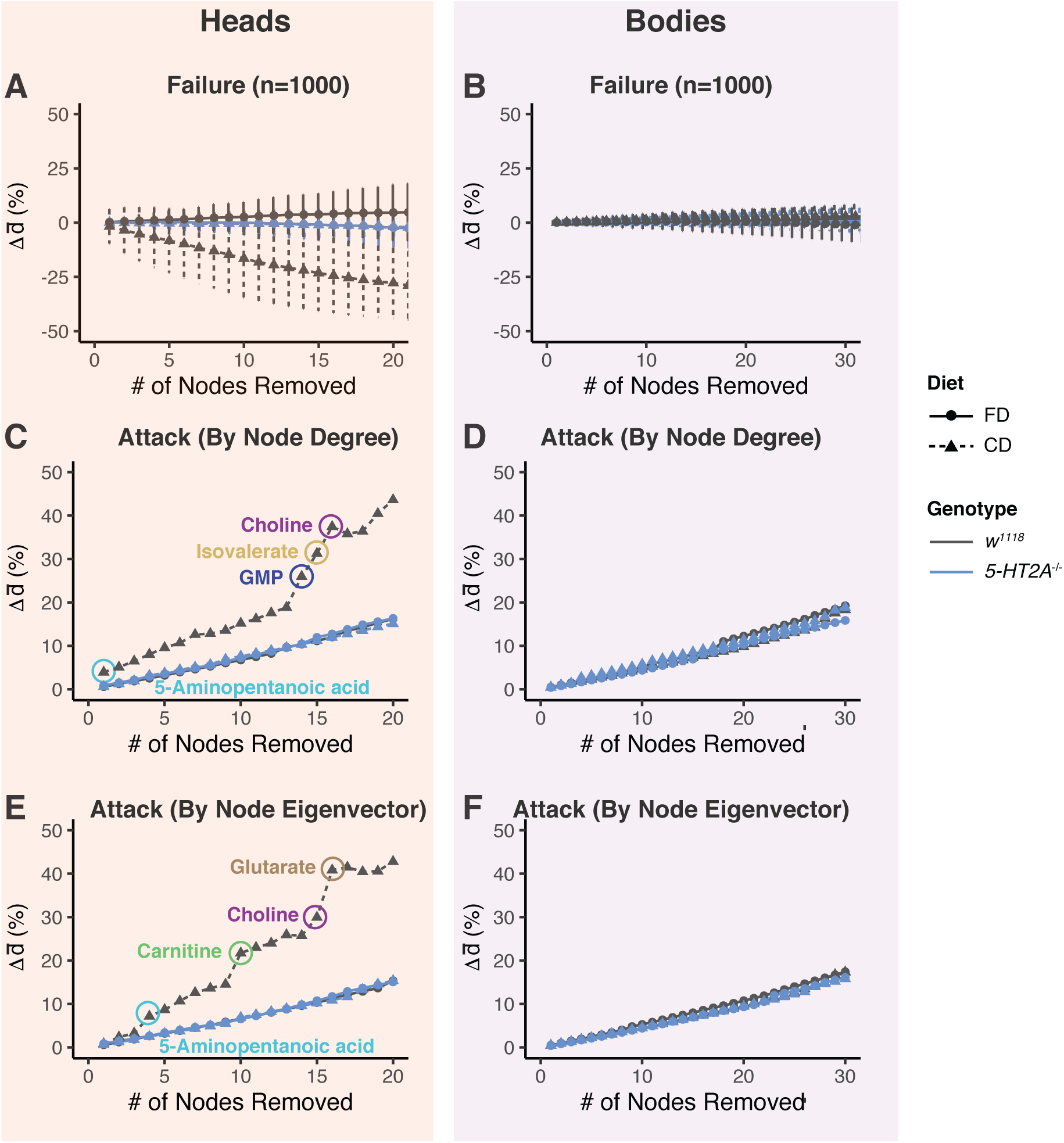
Serotonin signaling mediates the decrease in network robustness in response to nutrient choice. (**A-B**) Randomly removing nodes in the networks does not increase average shortest distance in heads (**A**) or bodies (**B**). Dots and error bars indicate the mean and standard deviation from 1,000 simulations respectively. (**C-F**) Sequential removals following node centrality (estimated from node degree (**C-D**) or node eigenvector (**E-F**)) increase network average shortest distance 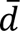 in both heads and bodies. Importantly, curves that represent the CD network in heads exhibit significant rises after removing several key metabolites, which are highlighted in circles.

### Dietary choice increases phenotypic vulnerability to genetic perturbations on the glutamine-**α**KG axis

Our systems analyses motivated us to test a working mechanistic model (Figure 6A) in which the effects of dietary choice on network structure, which include increased fragmentation as well as reduced connectivity, average shortest distance, and robustness, would increase organism-level vulnerability to genetic or environmental perturbations, akin to removal of network nodes, as measured by resistance to stress. Previously we demonstrated that the glutamine-αKG axis likely acts downstream of serotonin signaling to mediate the lifespan effects from dietary choice^31^. As a key TCA intermediate and a cofactor of DNA/histone demethylating enzymes, αKG sits at the intersection of mitochondrial metabolism and epigenetic regulation of aging. Manipulating this hub metabolite presumably influences multiple physiological pathways and might affect systematic robustness^36^.

**Figure 6.**
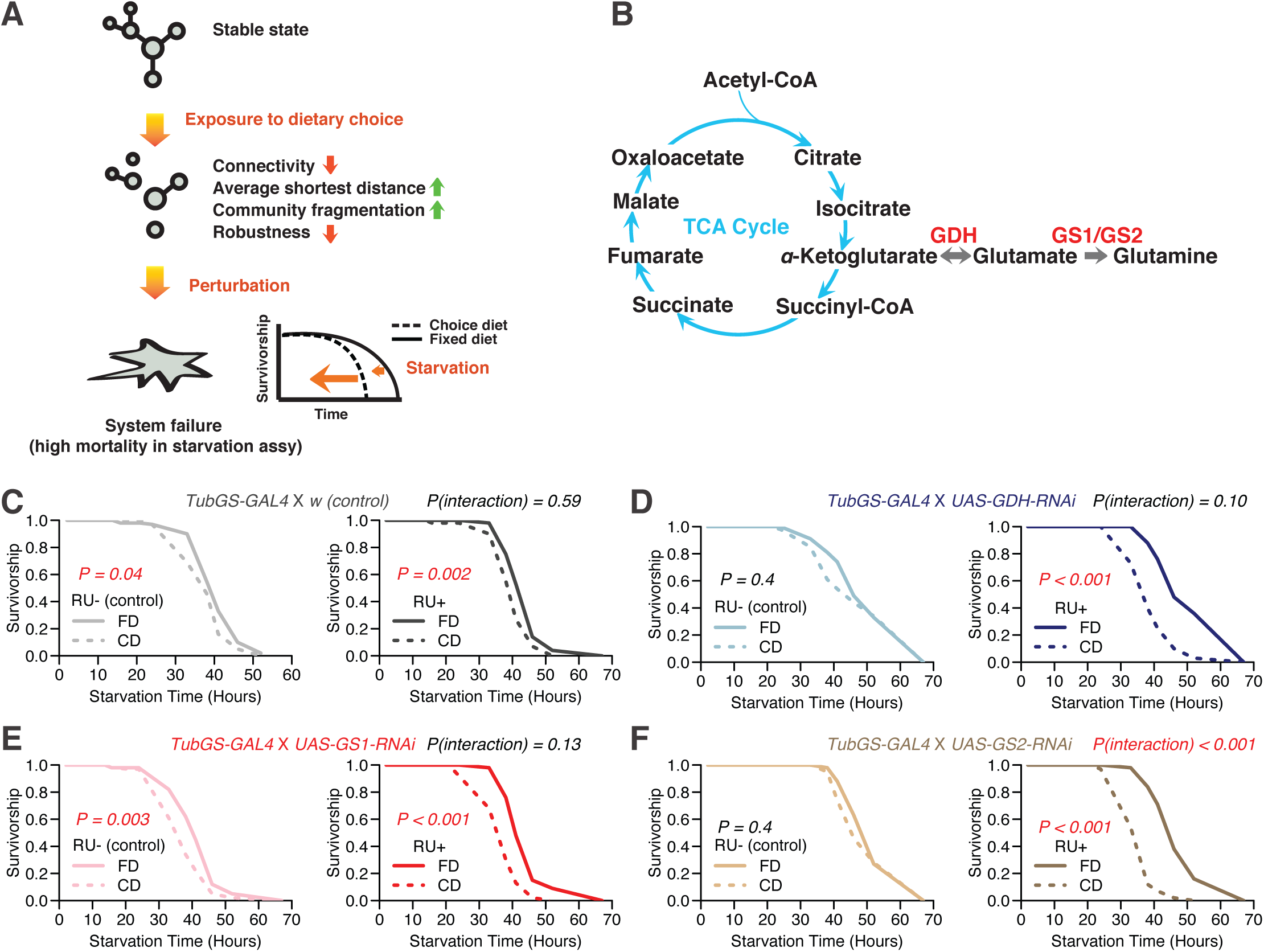
**Exposure to dietary choice increases vulnerability to genetic perturbations**. (**A**) Our working model. (**B**) glutamine-αKG axis plays important roles in fueling the TCA cycle, one of the central biochemical reactions in metabolic networks. (**C-F**) Upon starvation, knocking down key enzymes (*GDH*, *GS1* and *GS2*) on the glutamine-αKG axis exacerbates the dietary effects, suggesting flies that are exposed to a choice diet are more sensitive to stressors. *P*-values between dietary comparison are obtained by log- rank test. We also use cox-regression analysis to report *P*-value for the interaction term between genotypes and diets. Each treatment consists of six biological replicates, with 5-10 flies in each one. *P*- values from the interaction between diet (CD/FD) and genetic manipulation (RU+/RU-) are significant in the *GS2* knock-down group, and are marginal in the *GDH* or *GS1* knock-down group.

We therefore executed adult-specific, RNAi-mediated knock down of each of three key enzymes that control the glutamine-αKG axis: *GDH* (*Glutamate Dehydrogenase*), *GS1* (*Glutamine Synthetase 1*), and *GS2* (*Glutamine Synthetase 2*) (Figure 6B), and we measured the effect of diet on starvation resistance (see Methods for details). We predicted that the effect of knockdown on this phenotype would be magnified in flies fed a choice diet because of the putative fragility of the observed networks (Figure 6A). Consistent with our previous findings^31^, control males that were fed a choice diet for 10 days exhibited a smaller decline in starvation resistance relative to their siblings fed a fixed diet (Figure 6C). Importantly, knocking down any one of the three enzymes exacerbated the differences between a choice and a fixed diet, as predicted by our model (Figures 6D-6F). This effect was not caused by the transcriptional inducer, RU486, because control animals (*tub-GS-GAL4* coupled with *w*) responded as expected to dietary choice when RU486 was presented (Figure. 6C).

## Discussion

We discovered that nutrient presentation modulated fundamental network structures in metabolomes through serotonin signaling via the 2A receptor branch. We probed different aspects of network integrity, including: *i*) connectivity and average shortest distance, which revealed the degree to which the abundance of individual metabolites was responsive to the changes in the rest of the metabolomes; *ii*) community structure, which reflected the underlying organizations of the correlational networks and *iii*) robustness, which predicted how well the systems could handle extrinsic and intrinsic perturbation. Our analyses revealed that meal choice reshaped a highly interconnected metabolic network into a more fragmented one, which consisted of fewer edges and smaller communities, and that these structures were more vulnerable to perturbations. Importantly, all these effects were dependent to some extent on receptor 5-HT2A. While similar effects on metabolomes have been observed following dietary restriction^13^, our findings provide a molecular mechanism between nutrient sensing and network integrity, which, to our knowledge, are the first to do so.

We speculated that the changes on network structures through 5-HT2A were adaptive responses to a complex nutritional environment. Serotonin signaling has been shown to influence both energy states and behavior^25, 30^, and it seems plausible that 5-HT2A signaling is involved in coordinating a variety of responses to environmental nutrients in such a way as to maximize individual fitness. Although a less connected, more fragmented metabolome may be associated with a reduced lifespan^13^, it may be beneficial to overall fitness in a heterogenous environment, in which specific metabolic pathways might be responsive to individual nutrients. Improved metabolic efficiency and specialization in somatic cells are traits that are thought to be favored over longevity assurance^45, 46^.

We propose that the elevated mortality rates that flies experience when given a dietary choice may be due, at least in part, to a decrease in physiological robustness. A more fragmented, isolated network might be expected to increase organismal frailty and reduce resistance to other stressors, which was also indicated by our computational simulations (Figure 5). Our experimental results showed that the effects of genetic perturbation on starvation resistance were magnified when flies were given a dietary choice, which is consistent with this idea (Figure 6). This conjecture is also in line with previous findings that demonstrated a relationship between network integrity and aging in the transcriptomes and metabolomes of organisms across taxa, including nematodes^14^, fruit flies^13, 47^ and mice^11, 12, 48^. The consistency of this trend across evolutionarily distant species suggests that a decline of systematic robustness is an emergent property of aging. Its characteristics may offer useful aging biomarkers, which have proven elusive at the molecular level^49^.

Our study is among the few showing that network robustness and physiological robustness are associated and that systems-level adaptations to dietary conditions can be mediated by single genes, providing new insight into the mechanisms through which network integrity impacts aging. Knocking down critical metabolic enzymes was more deleterious in animals with vulnerable network structures (i.e., flies fed a choice diet, see Figure 6E-6F). On the other hand, loss of serotonin receptor 5-HT2A increased network robustness in a choice environment and also rescued choice-induced declines in starvation resistance^31^. Nevertheless, mechanisms that influence physiological robustness, as the manifestation of biological robustness occurs on many different levels^3^.

Going forward, a deeper understanding of the mechanisms through which serotonin signaling modulates lifespan may benefit from a focus on highly connected ‘hub’ molecules in the metabolic networks that we have identified. The nature of biological networks concentrates significant power in these hubs because they often govern major fluxes of metabolic information^34, 50^, and network hubs are often effective targets for interventions that accelerate or slow aging^16–18, 36^. At the molecular level, metabolic enzymes that sit at the branch point of metabolic reaction networks are considered more influential in these processes ^51^. Recent studies have exposed the identities of hub metabolites and their roles in modulating lifespan. Metabolites considered as the downstream integrators of nutrient metabolism and signaling pathways, including nicotinamide adenine dinucleotide [NAD+], reduced nicotinamide dinucleotide phosphate [NADH], α-ketoglutarate[αKG], and β-hydroxybutyrate [βHB], have emerged as key mediators of lifespan^36^. This is consistent with our observations that αKG appears to be one of the major effectors of serotonin signaling that modulates aging^31^ and physiology (Figure 5C-5F). The αKG/glutamine synthesis pathways may also be important in modulating the ability of diet restriction to increase lifespan^19^. More recently, it has been shown that the lifespan-extending effects of αKG are preserved in both flies^52^ and mice^53^. In the future, it is imperative to investigate the impact of these metabolites and metabolic pathways on network structures and metabolic communication and to better understand the mechanisms through which nutrient presentation influences lifespan.

## Materials and Methods

### Metabolomic Study

Metabolomics data were collected in a separate study^31^. We briefly summarized our methods here, while the full details can be found from our publication^31^. Males were exposed to the choice or fixed diet for 20 days (food was changed every 2-3 days) and then switched to the other diet (experimental groups) or the same diet (control groups). Forty to fifty heads or bodies were then homogenized for 20 sec in 200μl of a 1:4 (v:v) water : MeOH solvent mixture. Following the addition of 800 μl of methanol, the samples were incubated for 30 mins on dry ice, then homogenized again. The mixture was spun at 13000 RPM for 5 mins at 4°C, and the soluble extract was collected into vials. This extract was then dried in a speedvac at 30°C for approximately 3 hrs. Using a LC-QQQ-MS machine in the MRM mode, we targeted 205 metabolites in 25 important metabolic pathways, in both positive MS and negative MS modes. After removing internal controls and any metabolites missing from more than 8 out of 76 head samples (10.5%) and 8 out of 78 body samples (10.3%), we were left with 103 head metabolites and 125 body metabolites. Metabolite abundance for remaining missing values in this data set were log-transformed and imputed using the k-Nearest Neighbor (KNN) algorithm with the effects using the ComBat algorithm that implemented in the sva package of R Bioconductor. in then normalized the data to the standard normal distribution (μ=0, σ^2^=1).

### Correlation Network Analyses

Network analyses in this study were performed with the R language (version 4.0.0) within RStudio (version 1.2). Raw data and scripts are available as a GitHub repository (github.com/ylyu-fly/Metabolomics-FlyChoiceDiet).

#### Correlation analysis and network construction

To assess the relationship between metabolites for each biological condition, we estimated the significances of correlations using Spearman’s rank- order correlation coefficient test, which was computed from the abundance of metabolites within biological replicates (N=8-10). Afterwards, *P*-values were adjusted by FDR (Benjamini- Hochberg Procedure^54^), which we used for constructing adjacency matrices A = {*a*_*ij*_} and to infer the correlation network (non-directed graphs, as no directions were involved):

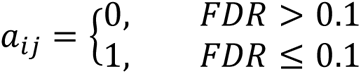

FDR = 0.1 was chosen as the cut-off followed standard practice. In correlation networks, nodes represented metabolites while edges represented links between them (where *a* =1). Visualization via the R package igraph^55^ revealed similar structures among the correlation networks across different diet and genotype groups. We observed a large, interonnected core group that represented the majority of the metabolites, which was accompanied by small groups as well as orphan metabolites. A pathway analysis using MetaboAnalystR^42^ was further applied to show the composition of the core group.

#### Permutation analysis

We generated permutated datasets to estimate the probability of observing the edge number differences between diets. We used the existing values with the group identify (i.e. genotype and diet) being shuffled for each metabolite. *P*-values were obtained from compare the real observation to 10,000 permutations.

#### Measures of network integrity

To investigate dietary influences on network integrity, we investigated four network attributes including:

(1) Connectivity (i.e. edge number/node degree);
(2) Average shortest distance (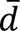);
(3) Community (module) structure, which revealed the underlying grouping pattern of correlation networks;
(4) Robustness, which assessed the influences of node removal on network attributes. We focused on the average shortest distance (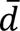), as it represented metabolic communication responsiveness which might impact aging56. Here we studied the robustness of network cores following our practice on the measures of average shortest distance.

#### Connectivity

We examined the dietary influences on network connectivity, specifically on the total edge numbers of the networks and on the number of neighbors for each node (i.e. degree). For total edge numbers, permutation testes were used to test whether the edge numbers in real networks were significantly deviated from those in random networks. Random networks were generated from 10,000 permutations; within each, group labels for each metabolite were shuffled independently. *P*-values were generated from the quantiles of the edge number (real experimental data versus the simulations). Similarly, *P*-values of the edge differences between diets (i.e. Δ_edge number_, defined as Nedge (CD) - Nedge(FD) in each realization) were calculated for both control and mutated flies. In addition to examining the dietary influences on the total edge number of networks, we also investigated the dietary effects on the frequencies of low-, intermediate- and high-degree nodes by plotting the distribution of edge number per node on a choice or a fixed diet.

#### Average shortest distance

Dietary effects on edge number suggested potential changes in how well metabolites connected to each other in these networks. To test this, we computed 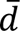, the average shortest distance between any two metabolites (in the core structures where every node was linked to each other) with the R package igraph^55^. As core groups are connected networks, there exists at least one path between any two metabolites in the group (i.e., a sequence of adjacent metabolites m0, m1, … mi, from m0 to mi each connected by at least one edge). The fewer steps along the shortest path from m0 to mi, the closer the two metabolites are considered to be.

#### Module analyses

We investigated the effects of meal choice on the community structure of the networks. Metabolites were separated into different modules, determined by the “leading eigenvector” approach^41^ with its implementation in igraph. The heart of this method is to obtain an optimized community classification by maximizing the modularity score (i.e. the number of edges within groups relative to that of random equivalent networks) across modules. Community structures were then visualized by the R package pheatmap. The choice-induced edge loss in the module was further quantified and visualized, through comparing the metabolite interactions between groups.

#### Network robustness analyses

We assessed the dietary influences on network robustness, with a focus on the effects of node removals on the average shortest distance (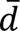). Following the practice of Albert *et al.*^43^, two approaches were used here to perturb network cores, where we either randomly removed nodes (denoted as network failure), or targeted the nodes that were considered as the ‘hubs’ of networks (denoted as network attack). For network failure, we calculated mean and standard deviation of Δ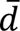 (changes in average shortest distance) from 1,000 simulations. For network attacks, two removing sequences were applied: we determined the centrality of nodes either by their degree or by their eigenvalue, with the node with the highest centrality (out of all the remaining ones) being removed first. The effects of sequential removal removal.

### Genetic Perturbation Analysis

#### Fly stock and husbandry

The laboratory stocks *w^11^*^18^, *UAS-GDH-RNAi* (BDSC#51473), *UAS-GS1-RNAi* (BDSC#40836) and *UAS-GS2-RNAi* (BDSC#40949) were purchased from the Bloomington *Drosophila* Resource Center. *Tub-GS-GAL4* flies were graciously provided by R. Davis (The Scripps Research Institute, Jupiter, FL). All fly stocks were maintained on a standard cornmeal-based larval growth medium (produced by LabScientific Inc. and purchased from Fisher Scientific) and in a controlled environment (25°C, 60% humidity) with a 12:12 hr Light:Dark cycle. We controlled the developmental larval density by manually aliquoting 32 uL of collected eggs into individual bottles containing 25 ml of food. Following eclosion, mixed-sex flies were kept on SY10 (10% (w/v) sucrose and 10% (w/v) yeast) medium for 2-3 days until they were used for experiments. Pioneer table sugar (purchased from Gordon Food Service, MI) and MP Biomedicals^TM^ Brewer’s Yeast (purchased from Fisher Sci.) were used through our study.

#### Dietary environments

To study the effects of dietary choice, we created food wells that were divided in the middle (Fig. 1A). This allowed us to expose the flies to two separate sources of food simultaneously. For these experiments, we either loaded different foods on each side (choice diet) or the same food on both sides(fixed diet). The choice diets contained 10% (w/v) sucrose or 10% (w/v) yeast on either side, and the fixed diets contained a mix of 10% (w/v) sucrose and 10% (w/v) yeast on each side.

#### Drug administration

We knocked down metabolic enzymes using the GeneSwitch system^57^.

The transcriptional inducer RU486 (mifepristone) was purchased from Sigma-Aldrich. Drug was first dissolved in 80% (v/v) ethanol at 10 mM concentration, marked by blue dye (5% (w/v) FD&C Blue No. 1; Spectrum Chemical), and stored at -20°C. For experimental food, 200 uM RU486 or the same dilution of control vehicle (80% (v/v) ethanol) was made from the stock and added to the food.

#### Starvation assay

We used a starvation survival assay to test the effects of genetic perturbations on organismal robustness. Once-mated, 2-3 days old male flies were kept on experimental (RU+) or control (RU-) food for 10 days (food was changed every 2-3 days) to activate the GeneSwitch system^57^. Afterward, we transferred flies to fresh vials containing 1% (w/v) agar. The number of dead flies was recorded approximately every 2-5 hrs using the DLife system^58^. Throughout the entire experiment flies were kept in constant temperature (25°C) and humidity (60%) conditions with a 12:12 hr Light:Dark cycle.

## Statistics

Fisher’s exact test was applied to analyze the effects of diet and genotype on the number of highly correlated metabolite pairs (defined as |*ρ*|≥ 0.8) against that of the rest pairs (i.e. |*ρ*|<0.8). Permutation analyses were used to assess the dietary influences on the number of edges in the correlational networks. To test the effects of diet and genotype involved in average shortest distance (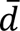), we performed two-way ANOVA (on the log-transformed 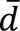). In the starvation resistance assay, pairwise comparisons between different treatment survivorship curves were carried out using the statistical package R within DLife^58^. For testing the interaction between genotypes and diets, we used Cox-regression analysis^59^ to estimate *P*-value for the interaction term.

## Data and Code Availability

Raw data, results and scripts that were used for this study can be obtained from our GitHub repository (github.com/ylyu-fly/Metabolomics-FlyChoiceDiet).

## Acknowledgements

We would like to thank Kelly Jin from the Promislow laboratory for her help with constructing correlational networks. This research was supported by the US National Institutes of Health, National Institute on Aging (R01 AG051649, R01 AG030593 to S.D.P. and R01 AG049494, R01 AG057330, and NSF grant DMS1561814 to D.E.L.P.), the Glenn Medical Foundation (to S.D.P.) and the Burroughs Wellcome Fund Collaborative Research Travel Grant (No. BWF1017452 to Y.L.).

## References

1. Barabasi, A. L. & Oltvai, Z. N. Network biology: understanding the cell’s functional organization. Nat Rev Genet 5, 101–113, doi:10.1038/nrg1272 (2004).

2. Hartwell, L. H., Hopfield, J. J., Leibler, S. & Murray, A. W. From molecular to modular cell biology. Nature 402, C47–52, doi:10.1038/35011540 (1999).

3. Kitano, H. Biological robustness. Nat Rev Genet 5, 826–837, doi:10.1038/nrg1471 (2004).

4. von Dassow, G., Meir, E., Munro, E. M. & Odell, G. M. The segment polarity network is a robust developmental module. Nature 406, 188–192, doi:10.1038/35018085 (2000).

5. Balaban, N. Q., Merrin, J., Chait, R., Kowalik, L. & Leibler, S. Bacterial persistence as a phenotypic switch. Science 305, 1622–1625, doi:10.1126/science.1099390 (2004).

6. Li, X., Cassidy, J. J., Reinke, C. A., Fischboeck, S. & Carthew, R. W. A microRNA imparts robustness against environmental fluctuation during development. Cell 137, 273–282, doi:10.1016/j.cell.2009.01.058 (2009).

7. Hou, L. et al. A systems approach to reverse engineer lifespan extension by dietary restriction. Cell Metab 23, 529–540, doi:10.1016/j.cmet.2016.02.002 (2016).

8. Riera, C. E., Merkwirth, C., De Magalhaes Filho, C. D. & Dillin, A. Signaling networks determining life span. Annu Rev Biochem 85, 35–64, doi:10.1146/annurev-biochem-060815-014451 (2016).

9. Soltow, Q. A., Jones, D. P. & Promislow, D. E. A network perspective on metabolism and aging. Integrative and comparative biology 50, 844–854, doi:10.1093/icb/icq094 (2010).

10. Hoffman, J. M., Lyu, Y., Pletcher, S. D. & Promislow, D. E. L. Proteomics and metabolomics in ageing research: from biomarkers to systems biology. Essays in biochemistry 61, 379–388, doi:10.1042/ebc20160083 (2017).

11. Bahar, R. et al. Increased cell-to-cell variation in gene expression in ageing mouse heart. Nature 441, 1011–1014, doi:10.1038/nature04844 (2006).

12. Southworth, L. K., Owen, A. B. & Kim, S. K. Aging mice show a decreasing correlation of gene expression within genetic modules. PLoS genetics 5, e1000776, doi:10.1371/journal.pgen.1000776 (2009).

13. Laye, M. J., Tran, V., Jones, D. P., Kapahi, P. & Promislow, D. E. The effects of age and dietary restriction on the tissue-specific metabolome of *Drosophila*. Aging Cell 14, 797–808, doi:10.1111/acel.12358 (2015).

14. Priebe, S. et al. Extension of life span by impaired glucose metabolism in *Caenorhabditis elegans* is accompanied by structural rearrangements of the transcriptomic network. PLoS One 8, e77776, doi:10.1371/journal.pone.0077776 (2013).

15. Promislow, D. E. Protein networks, pleiotropy and the evolution of senescence. Proc Biol Sci 271, 1225–1234, doi:10.1098/rspb.2004.2732 (2004).

16. Bell, R. et al. A human protein interaction network shows conservation of aging processes between human and invertebrate species. PLoS genetics 5, e1000414, doi:10.1371/journal.pgen.1000414 (2009).

17. Zhang, Q. et al. Systems-level analysis of human aging genes shed new light on mechanisms of aging. Human molecular genetics 25, 2934–2947, doi:10.1093/hmg/ddw145 (2016).

18. Xue, H. et al. A modular network model of aging. Mol Syst Biol 3, 147, doi:10.1038/msb4100189 (2007).

19. Jin, K. et al. Genetic and metabolomic architecture of variation in diet restriction- mediated lifespan extension in *Drosophila*. PLoS genetics 16, e1008835, doi:10.1371/journal.pgen.1008835 (2020).

20. Riera, C. E. & Dillin, A. Emerging role of sensory perception in aging and metabolism. Trends in endocrinology and metabolism: TEM 27, 294–303, doi:10.1016/j.tem.2016.03.007 (2016).

21. Gendron, C. M. et al. Neuronal mechanisms that drive organismal aging through the lens of perception. Annu Rev Physiol 82, 227–249, doi:10.1146/annurev-physiol-021119- 034440 (2020).

22. Wurtman, R. J. & Wurtman, J. J. Brain serotonin, carbohydrate-craving, obesity and depression. Obes Res 3 **Suppl 4**, 477S–480S, doi:10.1002/j.1550-8528.1995.tb00215.x (1995).

23. Olivier, B. Serotonin: a never-ending story. Eur J Pharmacol 753, 2–18, doi:10.1016/j.ejphar.2014.10.031 (2015).

24. Sze, J. Y., Victor, M., Loer, C., Shi, Y. & Ruvkun, G. Food and metabolic signalling defects in a *Caenorhabditis elegan*s serotonin-synthesis mutant. Nature 403, 560–564, doi:10.1038/35000609 (2000).

25. Srinivasan, S. et al. Serotonin regulates *C. elegans* fat and feeding through independent molecular mechanisms. Cell Metab 7, 533–544, doi:10.1016/j.cmet.2008.04.012 (2008).

26. Bouagnon, A. D. et al. Intestinal peroxisomal fatty acid β-oxidation regulates neural serotonin signaling through a feedback mechanism. PLoS biology 17, e3000242, doi:10.1371/journal.pbio.3000242 (2019).

27. Dierick, H. A. & Greenspan, R. J. Serotonin and neuropeptide F have opposite modulatory effects on fly aggression. Nat Genet 39, 678–682, doi:10.1038/ng2029 (2007).

28. Ries, A. S., Hermanns, T., Poeck, B. & Strauss, R. Serotonin modulates a depression-like state in *Drosophila* responsive to lithium treatment. Nature communications 8, 15738, doi:10.1038/ncomms15738 (2017).

29. Liu, C. et al. A serotonin-modulated circuit controls sleep architecture to regulate cognitive function independent of total sleep in *Drosophila*. Curr Biol 29, 3635–3646 e3635, doi:10.1016/j.cub.2019.08.079 (2019).

30. Albin, S. D. et al. A subset of serotonergic neurons evokes hunger in adult *Drosophila*. Curr Biol 25, 2435–2440, doi:10.1016/j.cub.2015.08.005 (2015).

31. Lyu, Y. et al. *Drosophila* serotonin 2A receptor signaling coordinates central metabolic processes to modulate aging in response to nutrient choice. Elife 10, e59399, doi:10.7554/eLife.59399 (2021).

32. Ro, J. et al. Serotonin signaling mediates protein valuation and aging. Elife 5, doi:10.7554/eLife.16843 (2016).

33. Nichols, C. D. 5-HT2 receptors in *Drosophila* are expressed in the brain and modulate aspects of circadian behaviors. Developmental neurobiology 67, 752–763, doi:10.1002/dneu.20370 (2007).

34. Barabasi, A. L. & Albert, R. Emergence of scaling in random networks. Science 286, 509–512 (1999).

35. Wagner, A. & Fell, D. A. The small world inside large metabolic networks. Proc Biol Sci 268, 1803–1810, doi:10.1098/rspb.2001.1711 (2001).

36. Sharma, R. & Ramanathan, A. The aging metabolome-biomarkers to hub metabolites. Proteomics 20, e1800407, doi:10.1002/pmic.201800407 (2020).

37. Kolaczyk, E. D. & Csárdi, G. Statistical analysis of network data with R. Vol. 65 (Springer, 2014).

38. Milgram, S. The small world problem. Psychology today 2, 60–67 (1967).

39. de Sola Pool, I. & Kochen, M. Contacts and influence. Soc Netw 1, 5–51 (1978).

40. Girvan, M. & Newman, M. E. Community structure in social and biological networks. Proc Natl Acad Sci U S A 99, 7821–7826, doi:10.1073/pnas.122653799 (2002).

41. Newman, M. E. Modularity and community structure in networks. Proc Natl Acad Sci U S A 103, 8577–8582, doi:10.1073/pnas.0601602103 (2006).

42. Pang, Z., Chong, J., Li, S. & Xia, J. MetaboAnalystR 3.0: Toward an optimized workflow for global metabolomics. Metabolites 10, doi:10.3390/metabo10050186 (2020).

43. Albert, R., Jeong, H. & Barabasi, A. L. Error and attack tolerance of complex networks. Nature 406, 378–382, doi:10.1038/35019019 (2000).

44. Iyer, S., Killingback, T., Sundaram, B. & Wang, Z. Attack robustness and centrality of complex networks. PLoS One 8, e59613, doi:10.1371/journal.pone.0059613 (2013).

45. Kowald, A. & Kirkwood, T. B. A network theory of ageing: the interactions of defective mitochondria, aberrant proteins, free radicals and scavengers in the ageing process. Mutation research 316, 209–236, doi:10.1016/s0921-8734(96)90005-3 (1996).

46. Kirkwood, T. B. Systems biology of ageing and longevity. Philos Trans R Soc Lond B Biol Sci 366, 64–70, doi:10.1098/rstb.2010.0275 (2011).

47. Avanesov, A. S. et al. Age- and diet-associated metabolome remodeling characterizes the aging process driven by damage accumulation. Elife 3, e02077, doi:10.7554/eLife.02077 (2014).

48. Derous, D. et al. The effects of graded levels of calorie restriction: VII. Topological rearrangement of hypothalamic aging networks. Aging (Albany NY) 8, 917–932, doi:10.18632/aging.100944 (2016).

49. Lopez-Otin, C., Blasco, M. A., Partridge, L., Serrano, M. & Kroemer, G. The hallmarks of aging. Cell 153, 1194–1217, doi:10.1016/j.cell.2013.05.039 (2013).

50. Jeong, H., Tombor, B., Albert, R., Oltvai, Z. N. & Barabasi, A. L. The large-scale organization of metabolic networks. Nature 407, 651–654, doi:10.1038/35036627 (2000).

51. Zhao, M. & Qu, H. Human liver rate-limiting enzymes influence metabolic flux via branch points and inhibitors. BMC Genomics 10 **Suppl 3**, S31, doi:10.1186/1471-2164-10-S3-S31 (2009).

52. Su, Y. et al. Alpha-ketoglutarate extends *Drosophila* lifespan by inhibiting mTOR and activating AMPK. Aging (Albany NY) 11, 4183–4197, doi:10.18632/aging.102045 (2019).

53. Asadi Shahmirzadi, A. et al. Alpha-ketoglutarate, an endogenous metabolite, extends lifespan and compresses morbidity in aging mice. Cell Metab 32, 447–456 e446, doi:10.1016/j.cmet.2020.08.004 (2020).

54. Benjamini, Y. & Hochberg, Y. Controlling the false discovery rate: a practical and powerful approach to multiple testing. J R Stat Soc Series B Stat Methodol 57, 289–300 (1995).

55. Csardi, G. & Nepusz, T. The igraph software package for complex network research. InterJournal, complex systems 1695, 1–9 (2006).

56. Smith, H. J., Sharma, A. & Mair, W. B. Metabolic communication and healthy aging: where should we focus our energy? Dev Cell 54, 196–211, doi:10.1016/j.devcel.2020.06.011 (2020).

57. Roman, G., Endo, K., Zong, L. & Davis, R. L. P[Switch], a system for spatial and temporal control of gene expression in *Drosophila melanogaster*. Proc Natl Acad Sci U S A 98, 12602–12607, doi:10.1073/pnas.221303998 (2001).

58. Linford, N. J., Bilgir, C., Ro, J. & Pletcher, S. D. Measurement of lifespan in *Drosophila melanogaster*. Journal of visualized experiments : JoVE, doi:10.3791/50068 (2013).

59. Cox, D. R. & Oakes, D. Analysis of survival data. Vol. 21 (CRC press, 1984).

