## Supplementary Figures for "Serotonin signaling modulates aging-associated metabolic network integrity in response to nutrient choice"

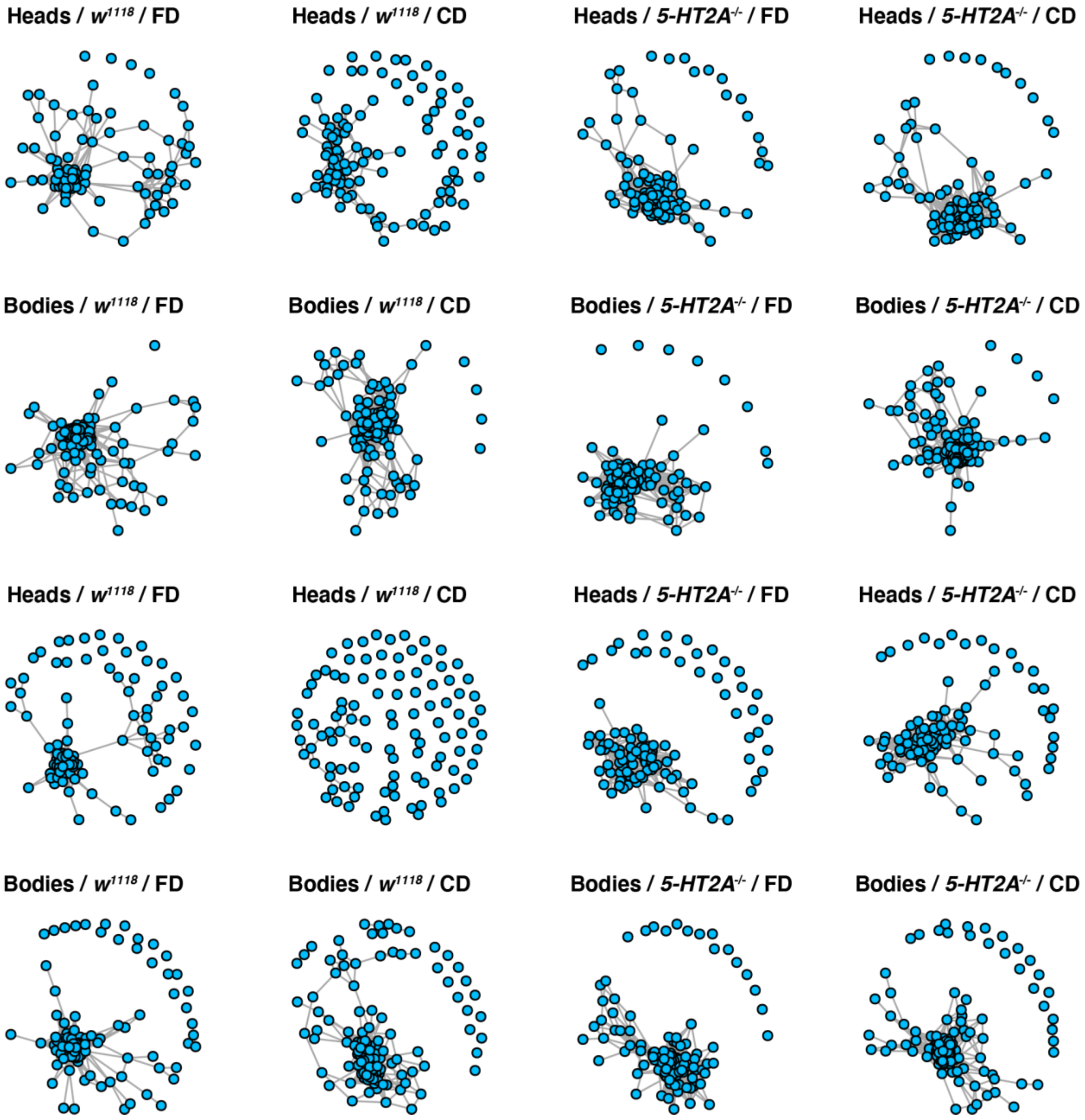


**Supplementary Figure 1. Network comparisons between two different FDR cut-offs**

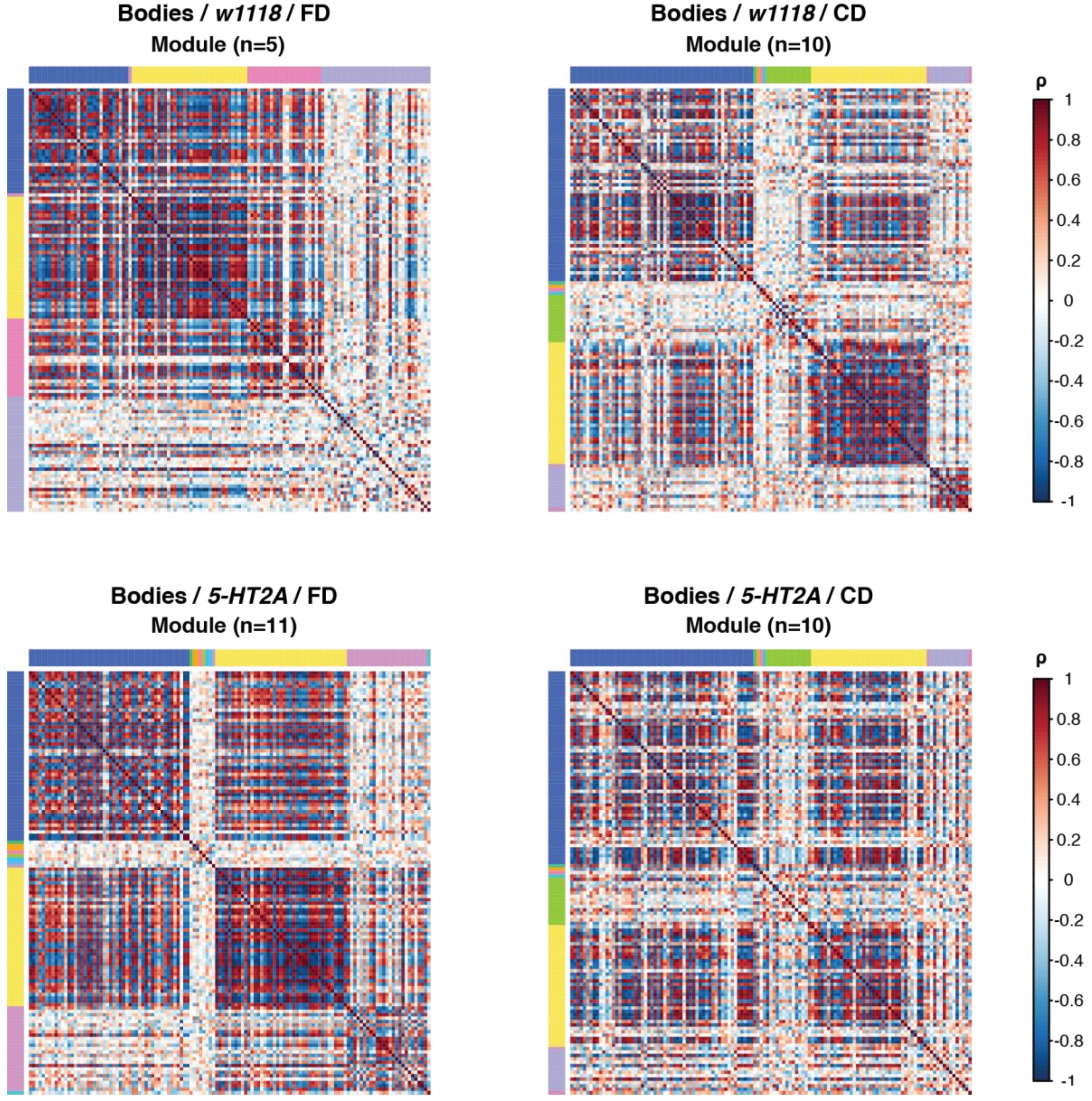


**Supplementary Figure 2. Choice-induced network fragmentation is not obvious in the bodies**
